# PCMD: A Multilevel Comparison Database of Intra- and Cross-species Metabolic Profiling in 530 Plant Species

**DOI:** 10.1101/2023.10.25.563765

**Authors:** Yue Hu, Yao Ruan, Xin-Le Zhao, Feng Jiang, Qiang Zhu, Qing-Ye Zhang, Qing-Yong Yang

**Affiliations:** National Key Laboratory of Crop Genetic Improvement, Hubei Hongshan Laboratory, Huazhong Agricultural University, Wuhan 430070, Hubei, China; Hubei Key Laboratory of Agricultural Bioinformatics, College of Informatics, Huazhong Agricultural University, Wuhan 430070, Hubei, China

**Keywords:** Comparative metabolomics, plant database, multi-level comparison, metabolite characteristics

## Abstract

Comparative metabolomics plays a crucial role in understanding gene function, exploring metabolite evolution, and improving crop genetic breeding. However, a systematic platform for comparing intra- and cross-species metabolites is currently lacking. In this study, we present the plant comparative metabolome database (PCMD; http://yanglab.hzau.edu.cn/PCMD), a comprehensive multi-level comparison database encompassing intra- and cross-species metabolic profiling in 530 plants. Remarkably, PCMD offers a multi-level platform for comparative metabolite analysis, allowing for the examination of metabolite characteristics across species at various levels including species, metabolites, pathways, and biological taxonomy. In addition, PCMD standardizes metabolite numbering, establishing a uniform system based on existing metabolite-related databases. The database also provides a range of user-friendly online tools, such as Species-comparison, Metabolites-enrichment, and ID conversion, enabling users to perform comparisons and enrichment analyses of metabolites across different species. PCMD stands out as the most comprehensive and species-rich comparative plant metabolomics database currently available, as demonstrated by two case studies that highlight its ability to supplement phylogenetic similarity mining among species through phylogenetic trees and offer new insights into the diversity and species-specificity of metabolites.

## Introduction

Plant metabolites are essential for the growth and development of plants (Shen et al., 2023). Comparing the variations in metabolites among different plants has immense benefits in unraveling their nutritional value, exploring metabolite evolution, and improving crop genetic breeding (Alseekh et al., 2021; Shen et al., 2023). A recent study indicated that flavonoids act as key metabolites underlying the evolutionary disparities between maize and rice (Deng et al., 2020). Additionally, a metabolomic analysis of three staple crops (wheat, rice, corn) and three fruits (grape, banana, mango) unveiled that crop-specific metabolites primarily contain flavonoids and lipids, whereas fruit-specific nutrients are predominantly flavonoids, providing metabolic evidence for a healthy human diet (Shi et al., 2022). Despite the existence of several databases facilitating research on plant metabolism, such as the Kyoto Encyclopedia of Genes and Genomes (KEGG) (Kanehisa and Goto, 2000), Plant Reactome (Naithani et al., 2020), and Plant Metabolic Network (PMN) (Hawkins et al., 2021), there remains a notable absence of a comprehensive platform for conducting comparative analyses of intra- and cross-species metabolites.

## Results

### Construction and overview of PCMD

In this study, we have constructed PCMD (Plant Comparative Metabolome Database), a multilevel comparison database for intra- and cross-species metabolic profiling (http://yanglab.hzau.edu.cn/PCMD) (**Fig. 1A**). PCMD serves as the most comprehensive and species-rich resource currently available in the field of comparative plant metabolomics, encompassing 530 plant species (**Supplemental Table 1**), 40,014 metabolites, 3,667 exogenous metabolites, 8,384 enzymes, 8,678 reactions, 14,521 experimentally-supported reactions and metabolites, and 39,964 literature (**Fig. 1B**). Notably, PCMD offers extensive species classification information, including taxonomy, reproductive characteristics, seed and leaf characteristics, and domestication information, enabling researchers to compare metabolite characteristics within and across species, or even at different taxonomic levels (**Fig. 1C, Supplemental Table 2**).

**Fig 1.**
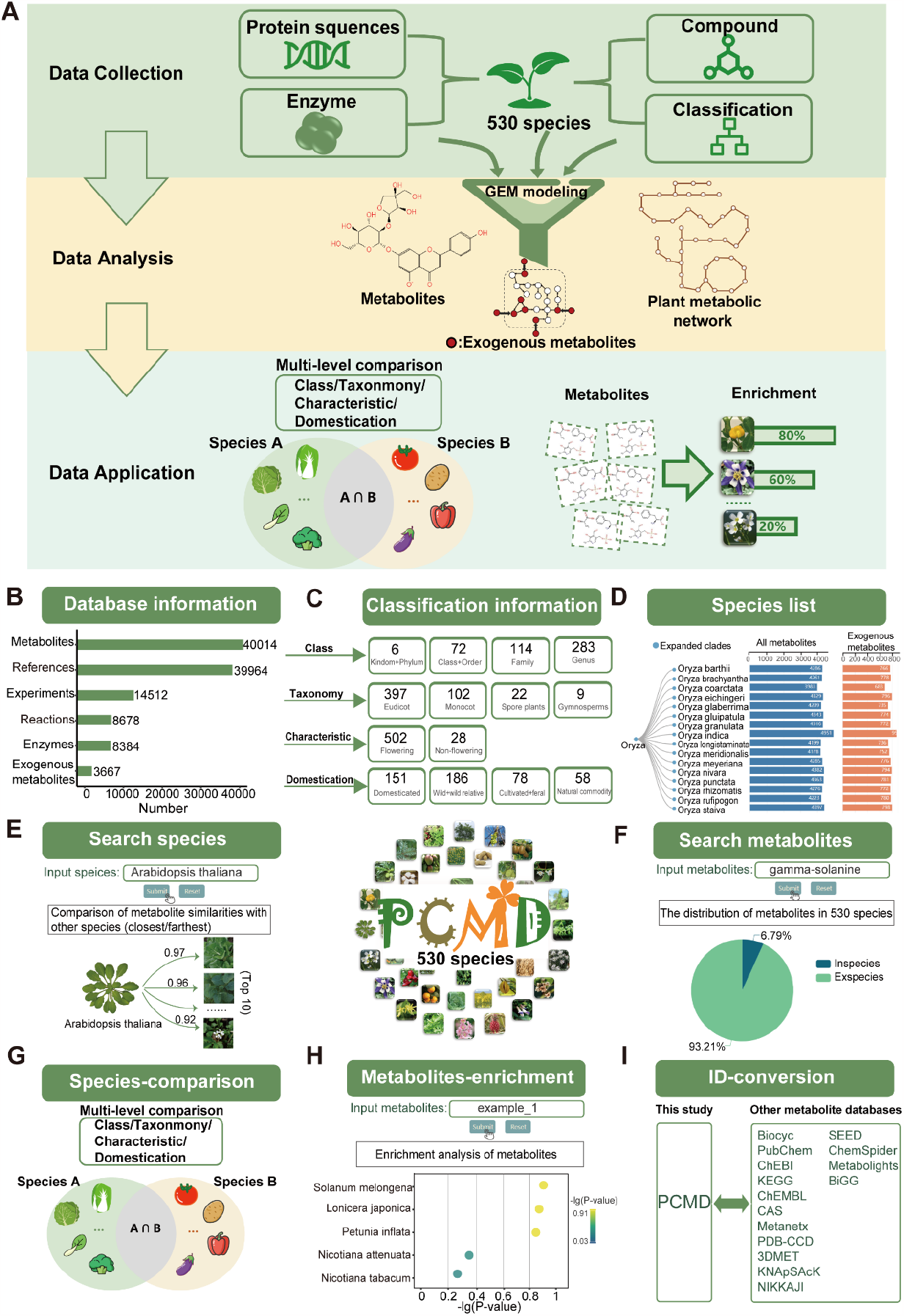
The architecture and representative functional modules of PCMD. (A) Workflow of PCMD. (B) Overview of the database information. (C) Classification information of 530 plant species. (D) The interface of Species list. (E) The interface of Search species. (F) The interface of Search metabolites. (G) Species-comparison tool. (H) Metabolites-enrichment tool. (I) ID-conversion tool.

To meet the varying needs of users in terms of retrieval and analysis, PCMD has meticulously designed six distinct modules: species, metabolites, reactions, literature, download, and help. The species module allows users to search for species, examine metabolite differences between species, and access metabolite information. On the Species list page, PCMD offers information on the classification tree of species and their respective rankings in the database for each plant (**Fig. 1D**). When users select or search for a species, PCMD provides basic species information, a comparison of metabolite similarities with other species, and more (**Fig. 1E; Supplemental Fig. 1**). Moreover, the metabolites module enables users to browse and search for metabolites and their structure and molecular weight. On the Search metabolites page, for instance, users can enter “gamma-solanine” to obtain basic information about the metabolite, its distribution in various species, a comparison of its structural similarities with other metabolites, etc (**Fig. 1F; Supplemental Fig. 2**).

### A multilevel comparative metabolite analysis platform across species

PCMD offers a comprehensive platform for multi-level comparative metabolite analysis across species. By comparing metabolites among different plants, new insights into the composition and characteristics of plant metabolomes can be gained. For instance, it is intriguing to discover that the plant with the most similar metabolites to the important oil crop *Brassica napus* is not from the *Brassica* genus, which is phylogenetically closer, but rather *Thlaspi arvense*, a member of the genus *Thlaspis* (**Supplemental Fig. 3**). This finding aligns with previous studies indicating the high similarity between *Thlaspi arvense*, an emerging oilseed crop, and *Brassica napus* (Warwick et al., 2002). Thus, comparing metabolite similarity in different plants holds promise for uncovering novel metabolic characteristics and serves as a valuable complement to mining phylogenetic similarities based on phylogenetic trees. Moreover, PCMD provides user-friendly online tools, such as Species-comparison, Metabolites-enrichment, ID conversion. The Species-comparison tool allows users to compare the specificity and commonness of metabolites between two different groups of plants at multiple taxonomic levels (**Fig. 1G, Supplemental Fig. 4**). For example, when comparing the Solanum and *Brassica* genera, differences in metabolites highlight alpha-chaconine and alpha-solanine as specific to Solanum plants. Specifically, *Brassica* plants lack the pathway for converting 16,22,26-trihydroxycholesterol to solanidine, impairing the synthesis of alpha-chaconine and alpha-solanine (**Supplemental Fig. 5, Supplemental Table 3**). Additionally, the Metabolites-enrichment tool enables users to analyze the enrichment of metabolites (**Fig. 1H**). The results provide basic information and classification of metabolites, the distribution of input metabolites across 530 species, enrichment analysis, and relevant literature (**Supplemental Fig. 6**). Remarkably, PCMD employs a standardized numbering system for metabolites, facilitating comparisons and enrichment analyses across various species. For example, the ID conversion tool allows for the conversion of metabolites from multiple published metabolomics databases (**Fig. 1I**).

## Discussion

Compared to other well-known plant metabolites databases such as KEGG, PMN, and Plant Reactome, PCMD possesses distinct characteristics. Firstly, PCMD serves as the most comprehensive and species-rich plant metabolomics database currently available (**Supplemental Table 4**). Secondly, it is the pioneering plant database that offers valuable resources for exploring exogenous metabolites. While plants rely on exogenous metabolites from the environment to sustain their growth and metabolism (Borenstein et al., 2008), several existing plant metabolite databases disregard the significance of these exogenous metabolites (**Supplemental Table 4**). Thirdly, PCMD serves as the first platform enabling the comparison of metabolic differences across various biological classification levels among different plant species. Plant Reactome primarily focuses on comparing pathways solely between rice and other plant species based on rice gene-orthology (Naithani *et al*., 2020), while each plant in PMN has its own relatively independent metabolic network database (Hawkins *et al*., 2021), both of which offer only a limited number of comparative analyses on metabolic differences across species. Finally, PCMD provides an ID-conversion tool that facilitates researchers in conducting joint analyses across multiple databases, thereby enhancing the practicality and applicability of the data. While KEGG does offer somewhat similar functions, it is worth noting that PCMD has the potential to connect more than 15 commonly used metabolic databases.

In the future, PCMD will focus on the following aspects: 1) Expanding the range of species included in the database and incorporating additional experimentally validated metabolites, reactions, and pathway-related data. 2) Optimizing the upload module to provide users with the capability to upload their own experimental data for comparative metabolite enrichment analysis. 3) Introducing new tools that facilitate exploration and analysis. For instance, the development of a metabolic network visualization tool will enable users to investigate the intricate relationships between metabolites and genes.

In summary, PCMD offers a valuable database for future research on the comparison of intra- and cross-species metabolites, providing new insights into metabolic diversity and species specificity. Additionally, PCMD’s contribution extends to promoting the standardization of plant metabolite identification and exploring the functional similarities of metabolites across different species.

## Author Contributions

Q.-Y.Y. and Q.-Y.Z. conceived and supervised the study. Y.H., Y.R. X.-L.Z. and F.J. collected the datasets, performed the bioinformatics analysis, and constructed the database. Q.-Y.Y., Q.-Y.Z. and Q.Z. revised the manuscript.

## Acknowledgments

No conflict of interest is declared.

