## Supplemental Information for "PCMD: A Multilevel Comparison Database of Intra- and Cross-species Metabolic Profiling in 530 Plant Species"

### Supplemental Materials and Methods

#### Collection of plant genomes and classification

The Plant Chemical Metabolomics Database (PCMD) serve as a comprehensive resource, encompassing 530 plant species. The genomic resources for these species were painstakingly gathered from various reliable sources, including the Ensembl Plants database (version 51), recent plant genome assembly studies, and other reputable published plant databases (**Supplemental Table 1**). Additionally, a wealth of species classification information was meticulously compiled, incorporating taxonomy, reproductive characteristics, seed and leaf characteristics, as well as domestication information. This classification data was sourced from the esteemed National Center for Biotechnology Information (NCBI) and a study conducted by Marks et al.(Marks et al., 2021) (**Supplemental Table 2**).

#### Construction of plant metabolic network

A genome-scale metabolic model (GEM) is a computational simulation representing the entirety of metabolic reactions occurring within a cell (Thiele and Palsson, 2010). By leveraging whole-genome sequences, GEMs have the capability to predict the reactions and metabolites present in organisms. In this study, plant metabolites are predicted based on GEMs. The construction of these GEMs relies on compilation of biochemical knowledge derived from diverse sources. The metabolic network for each plant is assembled *de novo* using the automated modeling tool RAVEN 2.0 (Wang et al., 2018).

RAVEN utilizes the MetaCyc database (Wang *et al.*, 2018) and the KEGG database (Kanehisa et al., 2021) in the process of *de novo* metabolic network reconstruction. The MetaCyc-based reconstruction module facilitates the identification of species-specific reactions and metabolites by comparing the amino acid sequences of enzymes to those in the MetaCyc database. This comparison generates a draft model as a result. Additionally, the KEGG-based reconstruction module employs a Hidden Markov Model (HMM) trained on genes annotated in KEGG to assess protein sequence similarities for the target species.

In this study, the MetaCyc-based draft models were generated using the "getMetaCycModelForOrganism" function available in RAVEN. The function was executed with default parameters to construct the draft models based on the MetaCyc database. For the KEGG-based draft model, we used the "getKEGGModelForOrganism" function in RAVEN, which employs an HMM trained on eukaryotic sequences with 90% sequence identity to query the plant proteome. Subsequently, we integrated the KEGG-based and MetaCyc-based draft models using the "combineMetaCycKEGGModels" function provided by RAVEN. This integration approach allowed us to leverage the complementary information from both the MetaCyc and KEGG databases, thereby enhancing the comprehensiveness and accuracy of the final draft model.

The complete metabolic networks were exported in a user-friendly Excel format. The resulting combined model integrates both KEGG and MetaCyc identifiers for metabolites. Subsequently, we carefully reviewed the metabolite identifiers, extracting and assigning consistent new identifiers to ensure uniformity. To address any potential absence of reaction formulas for certain reactions, we thoroughly examined all reactions within the model. We manually corrected any missing formulas and assigned new identifiers to each reaction to maintain accuracy and completeness.

**Identification of exogenous metabolites**

Exogenous metabolites play a crucial role in the growth and development of organisms, as they are compounds that cannot be synthesized internally and must be obtained from the external environment (Borenstein et al., 2008; Pavankumar et al., 2021). To identify exogenous metabolites in plants, we adopted a systematic approach.

Firstly, we extracted all reactions from existing plant metabolic models, which were then visualized as directed graphs. In these graphs, nodes represent metabolites, and edges denote the flow from reactants to products. To decompose these graphs into Strongly Connected Components (SCCs), we utilized the "nx.strongly_connected_components" function from the NetworkX library (version1.11; available on the GitHub Repository). It is important to note that metabolites within an SCC are interconnected, indicating that they share at least one common pathway.

A strongly connected component (SCC) with no incoming connections (in-degree) but at least one outgoing connection (out-degree) is commonly referred to as a “source component”. Any metabolite found within these source components can potentially be classified as an exogenous metabolite. To ensure the validity of the data, we have established the following criteria: Primarily, only those source components that contain a maximum of five metabolites were considered, following the recommended guidelines by Borenstein et al. Furthermore, to maintain the integrity of our analysis and eliminate potential outliers, reactions were disregarded if the corresponding directed graph had fewer than 10 nodes, also in accordance with the same study (Borenstein *et al.*, 2008). Such reactions were specifically labeled as isolated reactions.

**Comparative analysis of metabolite similarity among different species**

The Jaccard similarity coefficient was used to quantify the similarity of metabolites between two species. This coefficient evaluates the dissimilarity between the two species by calculating the ratio of unique metabolites found in each species to the total metabolites present in both species. In this research, two distinct sets of metabolites were considered, labeled as set A and set B. The Jaccard similarity coefficient, denoted as $J$, can be represented by the equation: $J$:

$$J(A,B) = \frac{|A \cap B|}{|A \cup B|} = \frac{|A \cap B|}{|A| + |B| - |A \cap B|}$$

**Identification of plant-specific metabolites**

To identify plant-specific metabolites and compare them among different families, genera, or two groups of plants, we utilized hypergeometric tests. Firstly, we constructed a presence-absence matrix where each row represented a metabolite and each column represented a different plant. A value of 1 indicated the presence of a metabolite in a particular plant, while a value of 0 indicated its absence. By analyzing these presence-absence matrices, we determined the number of metabolites present in different plant classes.

Secondly, we conducted hypergeometric tests to assess the level of enrichment of specific metabolites in these plant classes. These tests generated p-values that indicated the extent of enrichment, with smaller p-values denoting higher levels of enrichment. The hypergeometric tests were performed using the scipy package (Virtanen et al., 2020) (version1.7.3) in the Python programming language.

**System architecture and software for database construction**

PCMD (http://yanglab.hzau.edu.cn/PCMD) have been meticulously constructed. ThinkPHP (v.5.0.24) is used as the framework, with PHP serving as the foundational language. In addition, jQuery (v.3.6.0) is employed as the JavaScript library, and the Apache website server is utilized for parsing the MySQL database. The visualization of the data is made possible by incorporating various plugins, such as Bootstrap (https://getbootstrap.com), DataTables (https://datatables.net), and Echarts (https://echarts.apache.org). These plugins enable the creation of a wide range of charts, including bubble charts, pie charts, heat maps, line charts, and network diagrams (**Supplemental Table 5**). The database is designed to deliver a user-friendly experience and can be accessed online at no cost. Furthermore, it is compatible with multiple browsers such as Chrome (recommended), Opera, Firefox, Windows Edge, and macOS Safari.

### Supplemental Figures

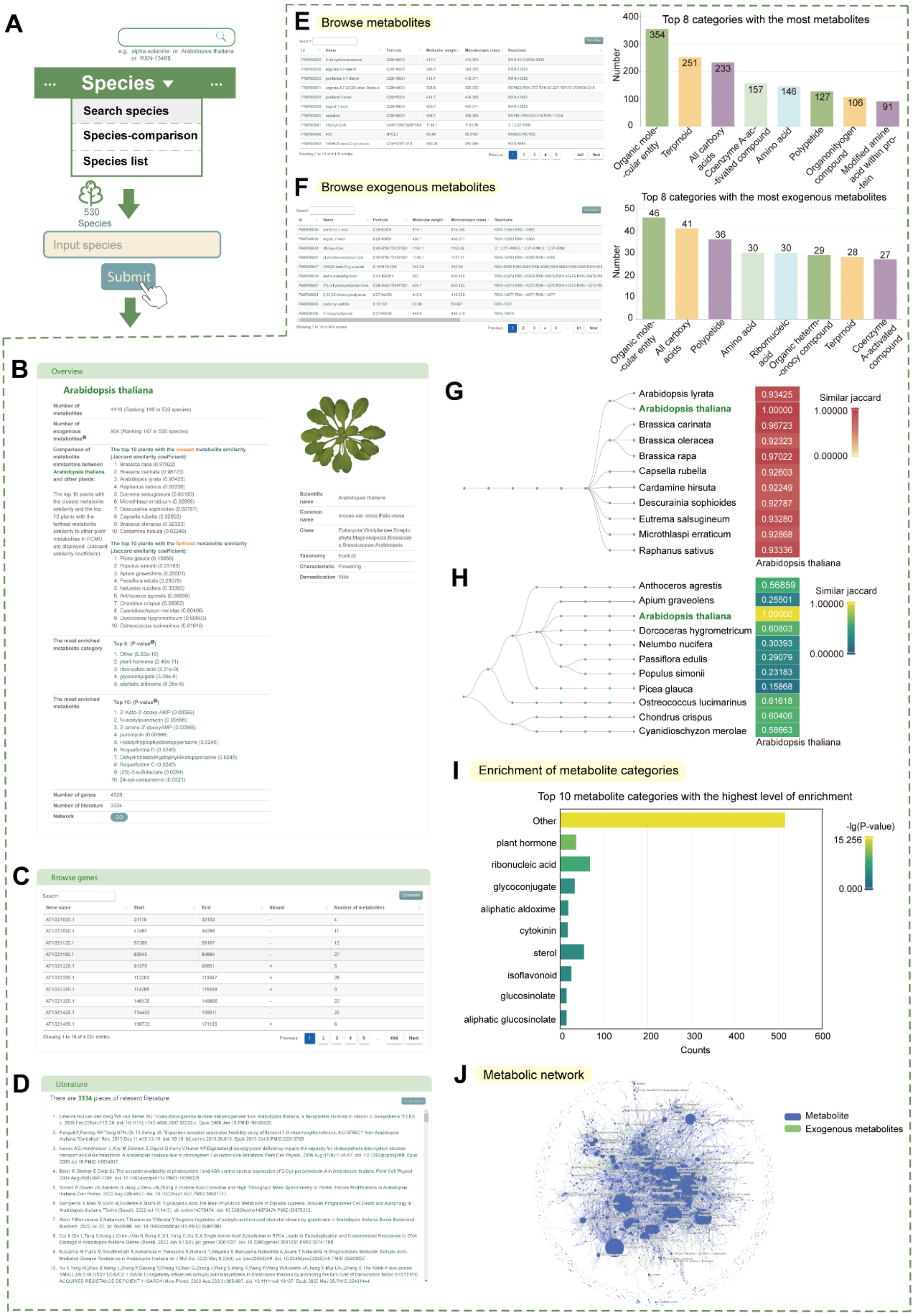


**Supplemental Fig. 1. Introduction to the "Search species" module.**

(A) Pipeline of the "Search species" module.

(B) Overview of the entered species.

(C) Results of genes associated with metabolites in the input species.
(D) List of literature related to the input species.
(E) Metabolites of the entered species and their classification.
(F) Exogenous metabolites of the entered species and their classification.
(G) Top 10 plants with the closest metabolite similarity to the input plant.
(H) Top 10 plants with the farthest metabolite similarity to the input plant.
(I) Bar plot ranking of the top 10 enrichment scores for metabolite categories with the highest level of enrichment.
(J) Metabolic network of the input species.

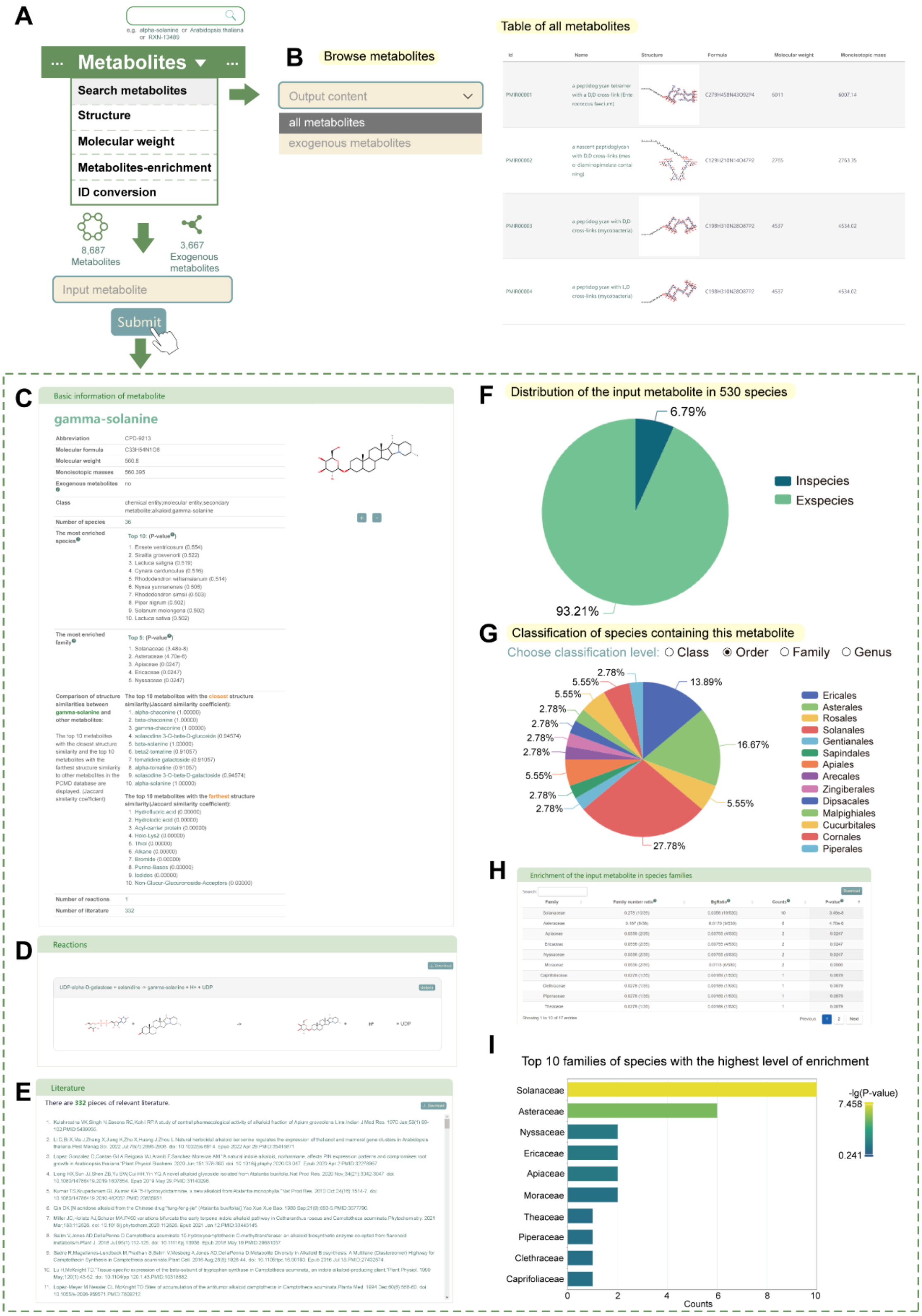


**Supplemental Fig. 2. Introduction to the "Search metabolites" module.**

(A) Pipeline of the "Search metabolites" module.

(B) Browse metabolites section, displaying basic information about metabolites in PCMD.

(C) Basic information of the input metabolite.

(D) Reactions associated with the input metabolite.
(E) List of literature related to the input metabolite.
(F) Distribution of the input metabolite in 530 species.
(G) Classification of species containing this metabolite, categorized by “Order”.
(H) Enrichment results of the input metabolite in species families.
(I) Bar plot ranking of the top 10 enrichment scores (–log10(P-value)) for species families with the input metabolite.

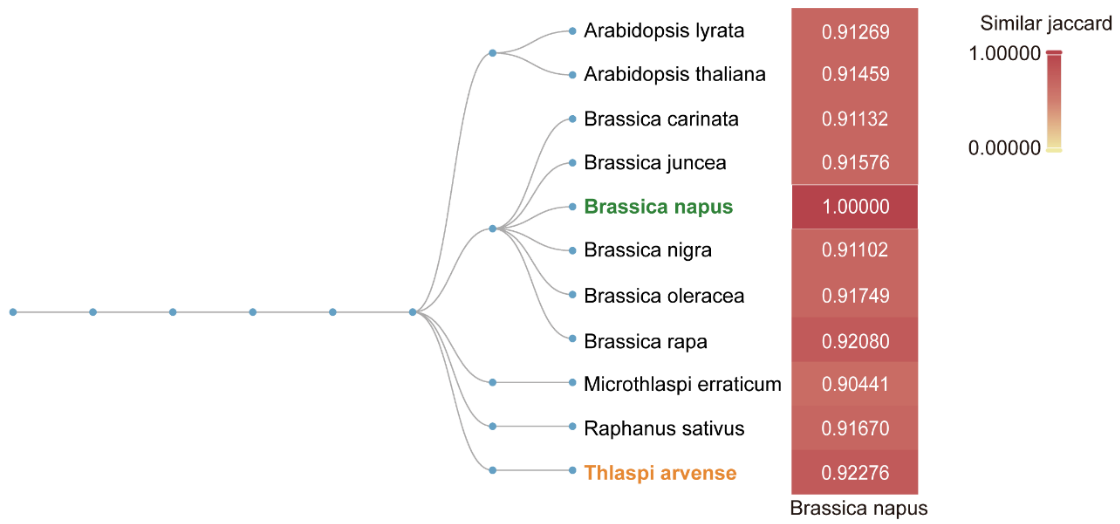


**Supplemental Fig. 3. Top 10 plants exhibiting the highest metabolite similarity to *Brassica napus* in PCMD analysis.**
*Brassica napus* is indicated by the green color, while *Thlaspi arvense* is highlighted in orange. The Jaccard similarity coefficient is used as the standardized measure for similarity.

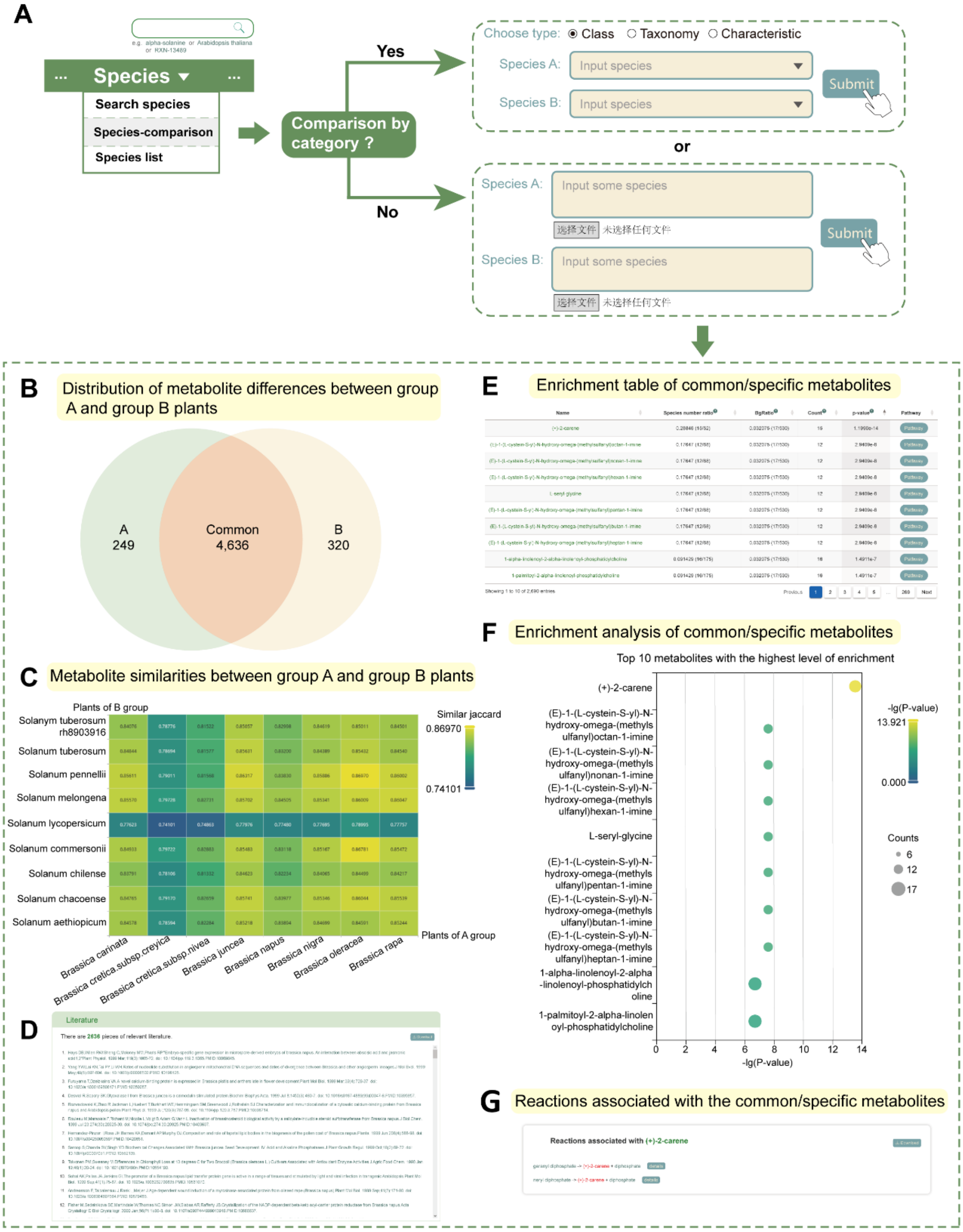


**Supplemental Fig. 4. Introduction to Species-comparison module.**

(A) Pipeline illustrating the comparison of metabolite differences between plants.

(B) Distribution of metabolite differences between group A and group B plants.

(C) Metabolite similarities between group A and group B plants.

(D) List of literature references on the metabolites in group A and group B plants in PCMD.

(E) Enrichment analysis results of common or specific metabolites between group A and group B plants.

(F) Enrichment analysis of common/specific metabolites between group A and group B plants. Bubble diagram depicting the top 10 metabolites with the highest level of enrichment.

(G) Reactions associated with the common or specific metabolites between group A and group B plants.

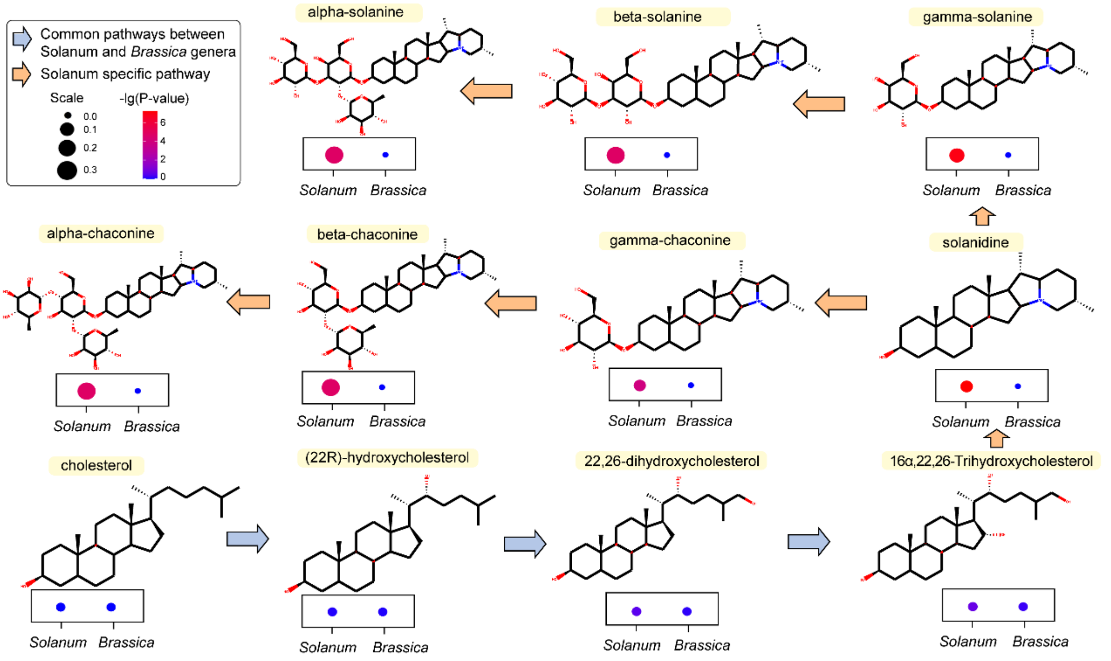


**Supplemental Fig. 5. Comparison of metabolites differences between Solanum and *Brassica* genera.**

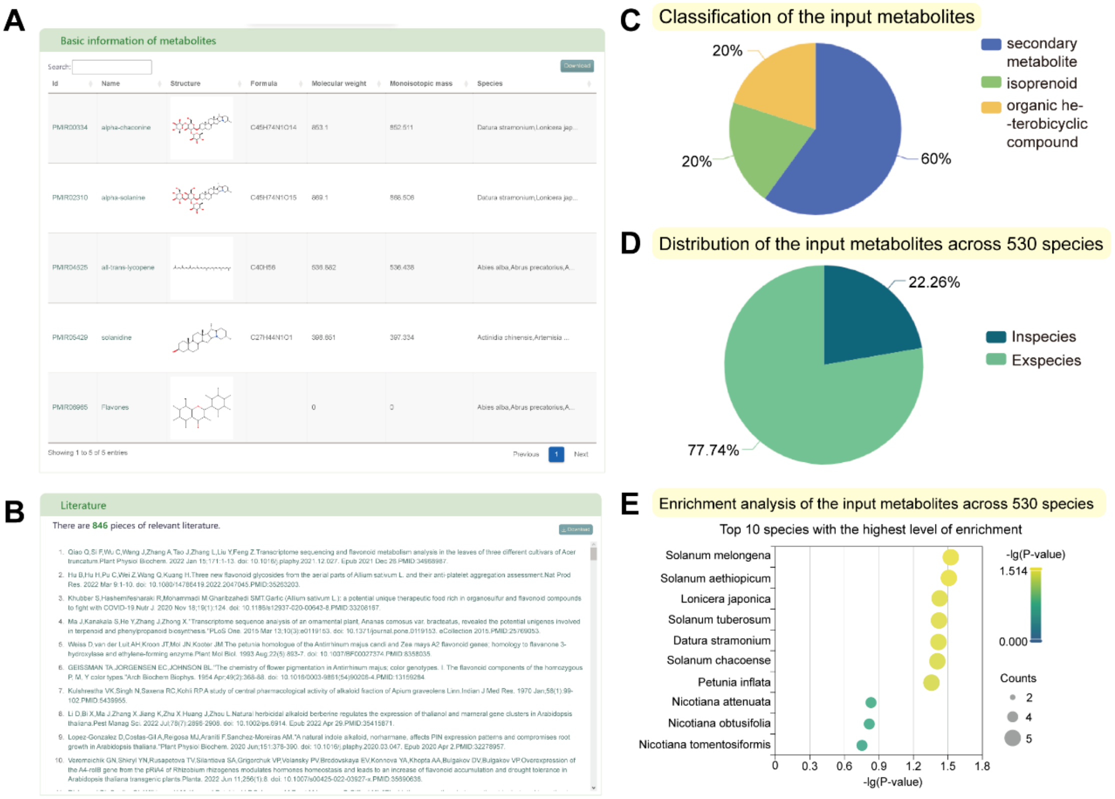


**Supplemental Fig. 6. Introduction to Metabolites-enrichment tool.**

(A) Basic information of the input metabolites.

(B) List of literature references related to the input metabolites.

(C) Classification of the input metabolites.

(D) Distribution of the input metabolites across 530 species.

(E) Enrichment analysis of the input metabolites across 530 species. The bubble diagram represents the top 10 species with the most significant enrichment.
